# cGAS integrates Toll-like receptor signaling in platelets to promote thromboinflammation

**DOI:** 10.64898/2026.11.02.763723

**Authors:** Amel Renaud Bidias, Laurence Arsenault, Samuel Fleury, Mélanie Welman, Mélissande Osouf, Marc Germain, Erwan Pernet, Marie Lordkipanidzé, Geneviève Pepin

## Abstract

Thromboinflammation is driven by innate immune signaling and promotes tissue injury during sepsis. Activation of platelets by bacterial products and alarmins amplifies inflammation, coagulation, and thrombotic dysfunction. However, the mechanisms by which these diverse inflammatory signals converge to elicit common platelet responses remain largely undefined. Mitochondria are key integrators of cellular stress responses and can activate innate immune pathways through the release of mitochondrial DNA and engagement of the DNA sensor cGAS. We therefore hypothesized that stimulation of innate immune receptors generates endogenous cGAS ligands within platelets, providing a unifying mechanism by which distinct inflammatory cues converge on cGAS-STING signaling. Here, we demonstrate that activation of TLR4, TLR1/2, and TLR5 engages a platelet-intrinsic mtDNA-cGAS-STING axis that promotes platelet activation. In a model of endotoxemia, genetic deletion of cGAS reduced platelet activation, thrombocytopenia, pyroptosis and the early systemic inflammatory response. Platelet transfer experiments further established that platelet-intrinsic cGAS is sufficient to amplify inflammatory responses *in vivo*. Mechanistically, we demonstrated that cytosolic accumulation of mtDNA and activation of cGAS-STING signaling pathway in platelets are dependent on mitochondrial calcium uniporter (MCU) and correlated with the loss of mitochondrial membrane potential and the production of mitochondrial reactive oxygen species. In contrast, canonical platelet agonists activated platelets independently of cGAS and failed to induce substantial cytosolic mtDNA accumulation. Importantly, this signaling architecture was conserved in human platelets, where TLR1/2-mediated activation required MCU- and STING-dependent signaling, whereas responses to thrombin receptor-activating peptide and collagen-related peptide remained largely unaffected. Together, these findings identify cGAS as an intracellular signal integration node that allows diverse innate immune receptors to converge on a common platelet activation pathway through sensing of mitochondria-derived DNA, allowing a rapid platelet response to microbial products.

## Introduction

Platelets are megakaryocyte-derived anucleate cells that are increasingly recognized as critical regulators of inflammation and thrombosis beyond their traditional role in hemostasis^1-3^. In addition to expressing canonical hemostatic receptors such as the collagen receptors (GPVI and GPIa/IIa) and the von Willebrand factor (VWF) receptors (GPIbα and GPIIb/IIIa), that are activated upon vessel injury to initiate coagulation, platelets possess a broad repertoire of innate immune receptors that enable detection of pathogen-associated molecular patterns (PAMPs) and danger-associated molecular patterns (DAMPs). Among platelet innate immune receptors, Toll-like receptors (TLRs) are major mediators of host responses to bacterial infection. Platelets express several cell membrane TLR family members, including TLR4, TLR1/2 and TLR5, which recognize lipopolysaccharide (LPS), bacterial lipoproteins and flagellin, respectively^3^.

These receptors allow platelets to actively contribute to thromboinflammatory responses during infection, tissue injury and systemic inflammatory diseases^1-13^. While the signaling pathways downstream of the receptors classically engaged during hemostasis are well described, our understanding of how innate immune receptors like TLRs contribute to platelet responses is still limited. In nucleated cells, activation of TLRs initiates multiple transcriptional and translational signaling programs that coordinate inflammatory responses^14,15^. However, platelets possess a limited capacity for *de novo* translation and therefore rely predominantly on post-translational signaling mechanisms to transform immune sensing into functional responses. Thus, how diverse innate immune receptors converge to promote platelet activation remains poorly understood.

The cyclic GMP-AMP synthase (cGAS)-stimulator of interferon genes (STING) pathway is a central component of innate immune surveillance^16,17^. cGAS detects cytosolic double-stranded DNA and catalyzes the synthesis of cyclic GMP-AMP (cGAMP), which activates STING and initiates inflammatory signaling cascades^18-20^. During infection, the cGAS-STING pathway is activated not only by microbial DNA but also by endogenous danger signals, particularly mitochondrial DNA (mtDNA) released into the cytosol following mitochondrial stress response^21-23^. Cytosolic accumulation of mtDNA has therefore emerged as a critical mechanism linking cellular stress to innate immune activation.

Recent studies have implicated cGAS-STING signaling in platelet-mediated thromboinflammatory responses^24-26^. Platelet STING activation promotes immunothrombosis and tissue injury during sepsis and myocardial infarction^24,25^. Currently it is suggested that platelet STING is activated through either the uptake of extracellular cGAMP generated by neighboring cells following cGAS activation in the vascular environment^24^ or through the internalization of dsDNA released by neutrophils during NET production^25^. An important limitation of current models is that they rely on extracellular cGAMP, and dsDNA supplied by neighboring cells, which would be expected to occur relatively late during inflammation. The existence of a platelet-intrinsic pathway capable of directly engaging cGAS would allow platelets to rapidly mobilize STING signaling during immune responses, without relying on the delayed accumulation of extracellular ligands generated by surrounding cells.

Emerging evidence suggests that platelet activation during severe inflammation is closely associated with mitochondrial depolarization^6,9,27-29^. Despite lacking a nucleus, platelets contain functional mitochondria that regulate metabolism, reactive oxygen species (ROS) production, calcium homeostasis and procoagulant activity^27^. Inflammatory stimuli such as LPS and the alarmin S100A8/A9 induce mitochondrial stress and mitochondrial depolarization in platelets^6,8^. Notably, these events have been linked to the NLRP3 inflammasome activation, platelet dysfunction and thrombocytopenia during sepsis^6,8^. We recently demonstrated that platelets express functional cGAS and that intracellular DNA delivery potentiates platelet activation through the cGAS-cGAMP-STING axis^26^. Together, these observations raise the possibility that mitochondrial depolarization generated during inflammatory stimulation may provide an endogenous source of DNA capable of activating platelet cGAS.

Because mitochondrial depolarization in response to calcium overload represents a common consequence of inflammatory receptor engagement^21,22^, we hypothesized that cGAS functions as an intracellular sensor of mtDNA generated downstream of innate immune stimulation. Such a mechanism would allow distinct innate immune receptors to converge on a common signaling pathway controlling platelet activation.

Here, we identify a platelet-intrinsic mtDNA-cGAS-STING signaling axis that links innate immune receptor stimulation to platelet activation. We demonstrate that activation of TLR4, TLR1/2 and TLR5 in platelets induces mitochondrial calcium uniporter (MCU)-dependent mitochondrial depolarization and cytosolic mtDNA accumulation, resulting in activation of cGAS-STING signaling. Functionally, this pathway can amplify platelet activation, platelet-leukocyte aggregate formation, inflammasome signaling, platelet procoagulant activity, and early inflammatory responses during endotoxemia. Using human platelets, we provide complementary evidence that TLR stimulation engages mitochondrial depolarization and STING signaling, supporting the translational relevance of this pathway. Our findings establish cGAS as a central integrator of inflammatory sensing in mouse and human platelets and reveal a mechanism through which diverse TLRs promote thromboinflammatory platelet responses.

## Results

### Platelet activation and early inflammatory responses to endotoxemia require cGAS

Endotoxemia triggers the rapid release of inflammatory mediators from multiple cell types, including platelets. To assess the various contributions of cGAS to systemic inflammation, circulating cytokines and platelet-derived inflammatory mediators were quantified following LPS challenge (10 mg /kg) by intraperitoneal of wild-type (WT) and cGAS knockout (*Cgas^-/-^*) mice. At 1 hour, IL-1β was undetectable in both genotypes, whereas plasma concentrations of CCL5, IL-6 and TNF-α were significantly reduced in *Cgas^-/-^* mice compared with wild-type (WT) animals (Fig. 1a, b, c, d). Notably, CCL5 is stored within platelet α-granules and rapidly released upon platelet activation^30,31^, suggesting that the reduction in circulating CCL5 may directly reflect impaired platelet activation in the absence of cGAS. By 3 hours, IL-1β increased and was modestly reduced in *Cgas^-/-^* mice (Fig. 1e). Because platelets have been reported to process and release IL-1β during inflammatory activation^32,33^ and to potentiate monocyte production of IL-1β^34^, this observation is consistent with a contribution of platelet cGAS to inflammatory signaling. In contrast, plasma concentrations of CCL5, IL-6 and TNF-α are similar in both genotypes at later time points (Fig. 1f, g, h), suggesting that cGAS primarily contributes to the early phase of the inflammatory response following endotoxin exposure.

**Fig. 1.**
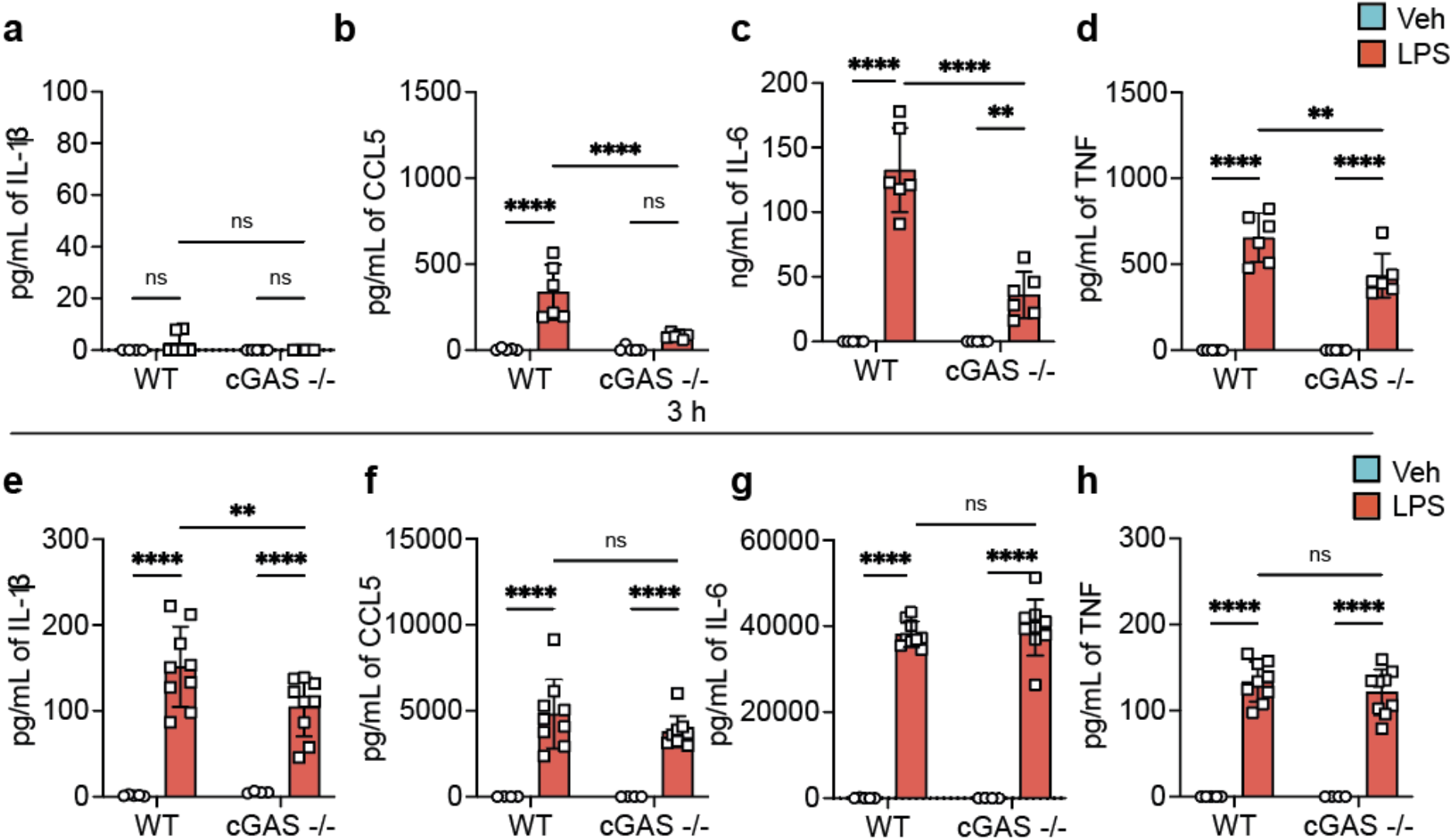
cGAS deficiency attenuates the early systemic inflammatory response to endotoxemia. **a-d**, Plasma concentrations of **a**, IL-1β, **b**, CCL5, **c**, IL-6 and **d**, TNF-α measured 1 hour following intraperitoneal injection of LPS (10 mg/kg) in WT and *Cgas*^-/-^ mice. **e-h**, Plasma concentrations of **e**, IL-1β, **f**, CCL5, **g**, IL-6 and **h**, TNF-α measured 3 hours following LPS challenge. Data are presented as mean ± SD. Each symbol represents one mouse. *n* = 4-9 biologically independent mice per group. Statistical significance was determined using two-way ANOVA followed by Tukey’s multiple-comparison test. *P < 0.01(**), P* < 0.0001 (****); ns, not significant.

Following endotoxemia, platelets are rapidly activated resulting in granule secretion and surface translocation of adhesion molecules like P-selectin (Cd62P), allowing the formation of platelet-leucocyte aggregates that rely on the interaction of P-selectin expressed by platelets and PSGL1 expressed by neutrophils and monocytes. One hour after endotoxin administration, platelet activation, measured by surface P-selectin expression, was significantly increased in WT but not in *Cgas^-/-^* mice (Fig. 2a). Consistent with this, platelet-neutrophil and platelet-monocyte-aggregate formation was also increased in WT mice at 1 hour post LPS (Fig. 2b, c). Although platelet activation remained elevated in WT but not *Cgas^-/-^* mice at 3 hours post-LPS, platelet-leukocyte aggregates returned to baseline levels (Supplementary Fig. 1). Of note, baseline levels were similar between genotypes suggesting a deficient response rather than a constitutive platelet phenotype.

**Fig. 2.**
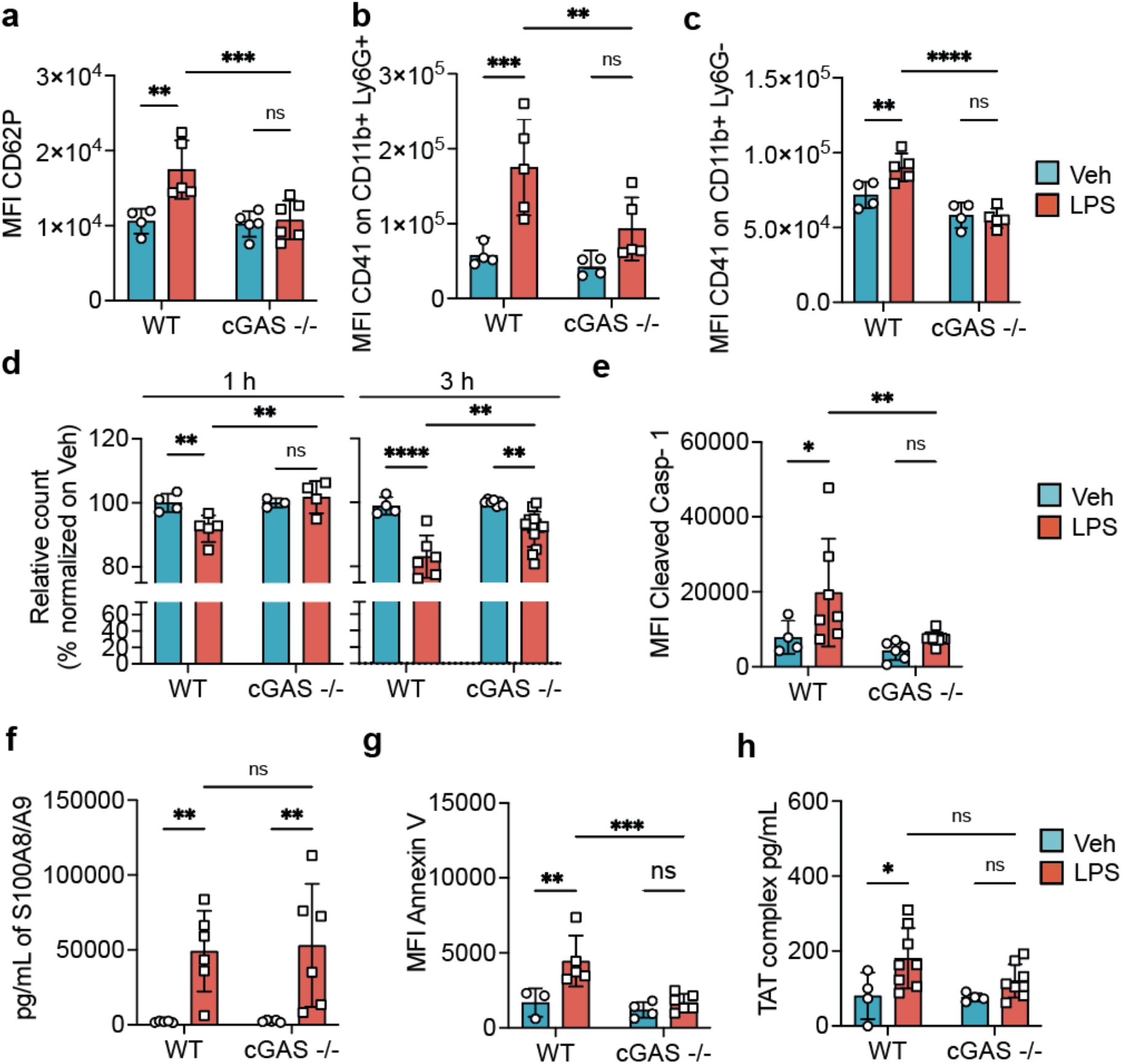
cGAS promotes platelet activation, inflammasome activation and procoagulant platelet formation during endotoxemia. **a-d**, Platelet responses measured following intraperitoneal injection of LPS (10 mg/kg) or sterile PBS in WT and *Cgas*^-/-^ mice. **a**, Platelet activation measured by MFI of surface CD62P expression on circulating CD41 platelets. **b**, Platelet-neutrophil aggregates quantified as CD41 fluorescence intensity on CD11b positive - Ly6G positive population. **c**, Platelet-monocytes aggregates quantified as CD41 fluorescence intensity on CD11b positive - Ly6G negative population. **d**, Relative platelet counts normalized to PBS-treated controls. **e-h**, Inflammatory platelet parameters measured 3 hours following LPS challenge. **e**, Cleaved caspase-1 measured by MFI of FLICA staining within platelet population. **f**, Plasma S100A8/A9 concentrations. **g**, Platelet phosphatidylserine exposure measured by Annexin V staining within platelet population. **h**, Plasma thrombin-antithrombin (TAT) complex concentrations. Data are presented as mean ± SD. Each symbol represents one mouse. *n* = 3-8 biologically independent mice per group. Statistical significance was determined using ordinary two-way ANOVA followed by Fisher’s least significant difference (LSD) multiple-comparisons test. *P < 0.05; **P < 0.01; ***P < 0.001; ****P < 0.0001; ns, not significant.

Thrombocytopenia during sepsis has been linked to both platelet consumption and shorten platelet lifespan and LPS challenge was reported to induce thrombocytopenia in mice^6,35,36^. When we measured platelet counts at 1 and 3 hours post-LPS injection, we observed a significant attenuation of platelet loss in *Cgas^-/-^* animals compared to WT (Fig. 2d). Because inflammatory signaling is a driver of platelet death pathways during endotoxemia and thereby contributes to thrombocytopenia, we investigated whether cGAS contributes to their reduction. Platelets can undergo different forms of cell death, including pyroptosis and apoptosis, characterized by NLRP3 inflammasome activation, caspase-1 cleavage, and IL-1β secretion^6,37,38^, and phosphatidylserine (PS) exposure^39,40^, respectively. Thus, we first assessed levels of cleaved caspase-1 in platelets (Fig. 2e) and found that in contrast to platelets from WT mice, pyroptosis was not induced in *Cgas^-/-^* mice. Platelet pyroptosis can be activated by the binding of the alarmin S100A8/A9, but circulating concentrations of S100A8/A9 measured three hours after LPS administration were comparable between genotypes (Fig. 2f), indicating that reduced pyroptosis in *Cgas^-/-^* mice was unlikely to result from altered systemic S100A8/A9 alarmin levels. Interestingly, PS exposure was also increased on platelets from WT mice during endotoxemia but not on *Cgas^-/-^* mice (Fig. 2g), confirming that cGAS contributes to platelet loss during endotoxemia. PS exposure is not only a hallmark of apoptotic platelets but is also a marker of the emergence of a procoagulant platelet phenotype that supports the assembly of the prothrombinase complex and thrombin generation. Consistent with the reduced PS exposure observed in *Cgas^-/-^* platelets, circulating thrombin-antithrombin (TAT) complex plasma levels increased in WT mice but not in *Cgas^-/-^* mice three hours after LPS administration (Fig. 2h), indicating that cGAS promotes both platelet death and the associated procoagulant response during endotoxemia.

### Platelet-intrinsic cGAS amplifies platelet activation and inflammatory responses during endotoxemia

To determine the specific contribution of platelet cGAS and exclude cGAS activity in other cellular compartments involved in host responses to endotoxin, we conducted a platelet transfusion experiment using *Cgas^-/-^*mice as the recipient to completely prevent cGAMP production by non-platelet cells. Endogenous platelets were depleted in *Cgas^-/-^*recipient mice by intraperitoneal injection of an anti-thrombocyte solution and animals were reconstituted with either WT or *Cgas^-/-^* platelets 12 hours post injection, before LPS challenge^41^. Transfer of WT platelets was sufficient to increase plasma concentrations of CCL5, TNF-α and IL-6 compared with transfer of *Cgas^-/-^*platelets (Fig. 3a, b, c), while plasma levels of the alarmins were not statistically different (Fig. 3d). These findings demonstrate that platelet cGAS actively contributes to the amplification of the systemic inflammatory response. Supporting a platelet origin for CCL5 release following endogenous cGAS activation, mice receiving WT platelets exhibited significantly greater platelet activation as assessed by CD62P surface expression one hour after LPS administration (Fig. 3e). Whereas platelet activation was clearly influenced by platelet cGAS, platelet-neutrophil aggregates, PS exposure and TAT complexes remained near baseline levels in both groups (Fig. 3f, g, h), suggesting that these responses may represent later events in the progression of endotoxemia.

**Fig. 3.**
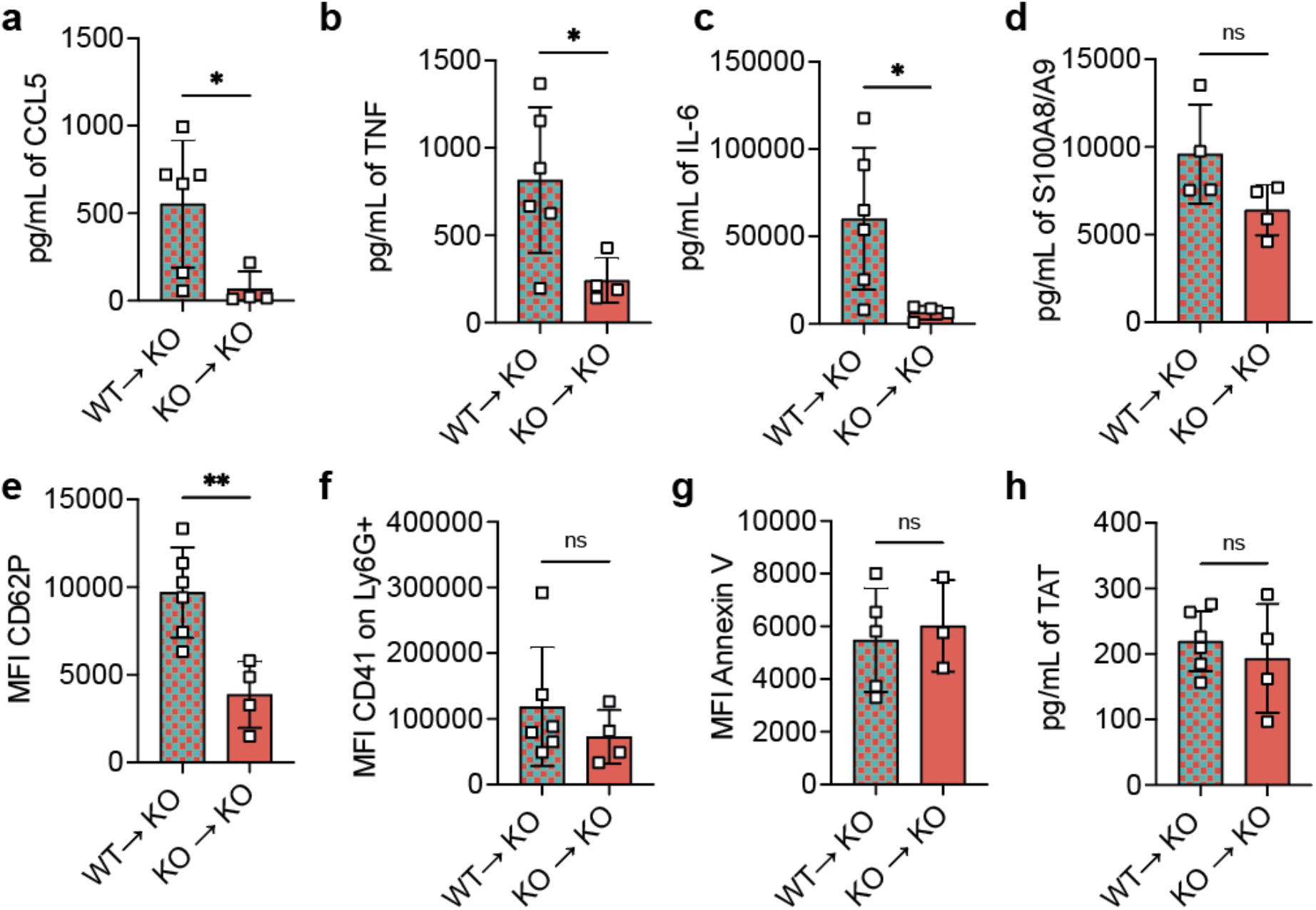
Platelet-intrinsic cGAS promotes platelet activation and inflammatory responses during endotoxemia. *Cgas*^-/-^ recipient mice were depleted of endogenous platelets and reconstituted with either donor WT or *Cgas*^-/-^ platelets prior to intraperitoneal LPS challenge (10 mg/kg). Plasma and platelet parameters were assessed 1 hour after LPS administration. **a-d**, Plasma concentrations of **a**, CCL5, **b**, TNF-α, **c**, IL-6 and **d**, S100A8/A9. **e**, Platelet activation quantified by surface CD62P expression on circulating platelets. **f**, Platelet-neutrophil aggregates quantified as CD41 mean fluorescence intensity on CD11b positive - Ly6G positive population. **g**, Platelet phosphatidylserine exposure measured by Annexin V staining. **f**, Plasma thrombin-antithrombin (TAT) complex concentrations. Data are presented as mean ± SD. Each symbol represents one transfused mouse. *n* = 4-6 biologically independent mice per group. Statistical analyses were performed using an ordinary two-tailed unpaired Mann-Whitney test. *P < 0.05; **P < 0.01; ns, not significant.

### TLR4-mediated platelet activation requires cGAS-STING signaling

Our *in vivo* analyses demonstrate that cGAS contributed to platelet activation, the formation of procoagulant platelets and caspase-1 activation during endotoxemia. To dissect the molecular mechanism leading to cGAS dependant platelet activity, isolated mouse platelets were exposed to LPS *in vitro*. LPS-induced platelet activation was significantly dependent on cGAS, as cGAS deficiency markedly prevented the increase of CD62P surface expression compared with WT controls (Fig. 4a). Interestingly, we observed that LPS was sufficient to promote an increase of PS exposure on WT but not on cGAS deficient platelets (Fig. 4b), suggesting that cGAS drives the formation of the prothrombinase complex. Stimulation with alarmin S100A8/A9 induced platelet activation in WT but not in cGAS deficient platelets (Fig. 4c). Pyroptosis, as measured by cleaved caspase-1 level and induced by the S100A8/A9 alarmin, was prevented in cGAS deficient platelets compared to WT (Fig. 4d). Similar results were obtained using the classical activator of pyroptosis LPS and nigericin (Supplementary fig. 2). To confirm the involvement of the downstream adaptor STING, WT platelets were stimulated with LPS or S100A8/A9 in the presence or absence of the STING inhibitor H-151^42^. Pharmacological inhibition of STING significantly reduced platelet activation and tended to reduce cleaved caspase-1 levels following stimulation with either agonist (Fig. 4e, f), confirming that STING functions downstream of cGAS in this pathway. These findings clearly establish endogenous cGAS signaling as a critical mediator linking inflammatory TLR4 stimulation to platelet activation and pyroptosis.

**Fig. 4.**
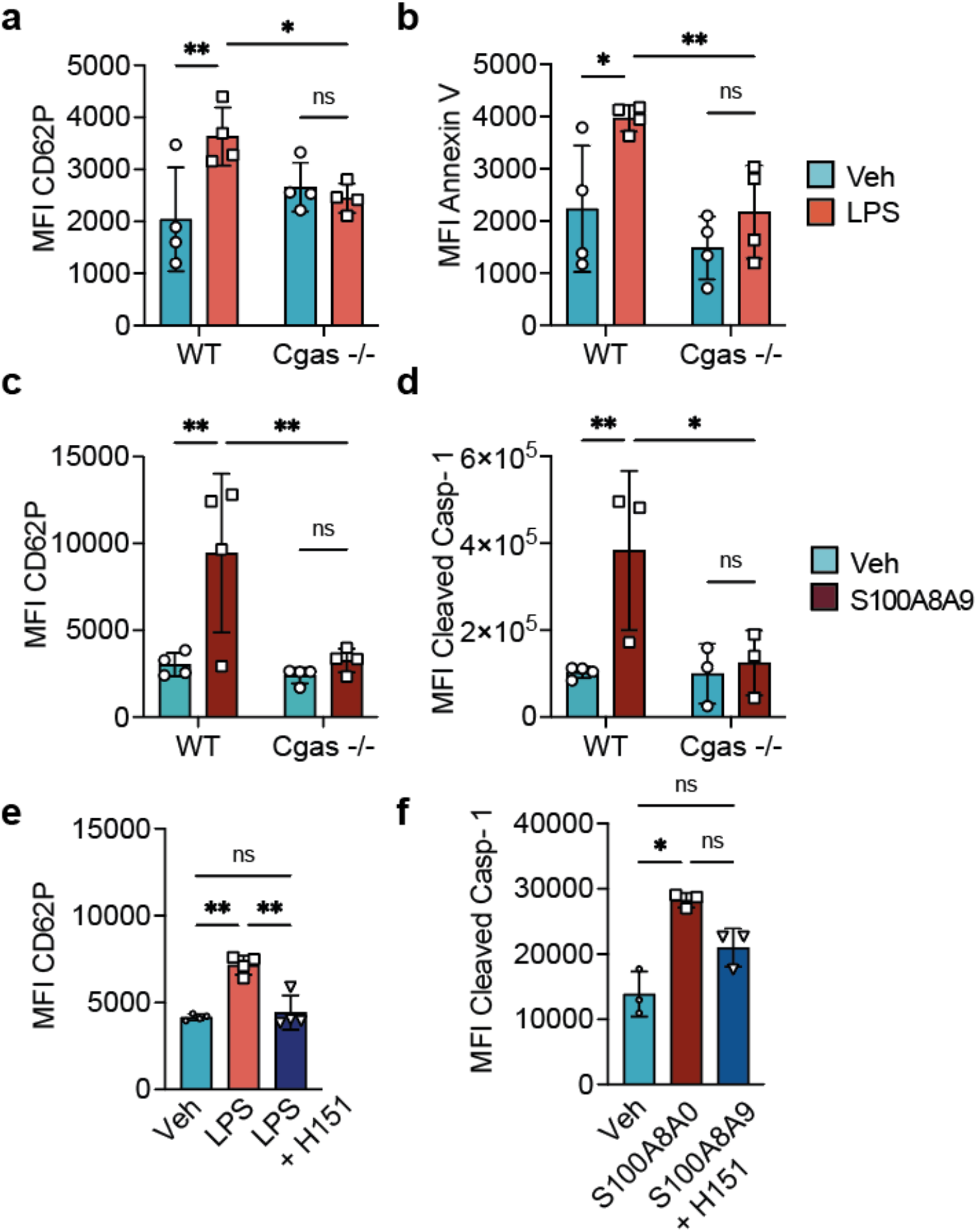
TLR4-mediated platelet activation requires cGAS-STING signaling Washed mouse platelets were stimulated *in vitro* as indicated. a,b Surface CD62P expression. (**a**) or phosphatidylserine exposure (**b**) on WT and *Cgas*^-/-^ platelets following stimulation with LPS. **c, d**, Surface CD62P expression on (**c**), or Cleaved Caspase-1 measured by FLICA mean fluorescence intensity in (**d**) WT and *Cgas*^-/-^ platelets following stimulation with S100A8/A9. **e** Surface CD62P expression on WT platelets stimulated with LPS and H151. **f** Cleaved Caspase-1 measured by FLICA mean fluorescence intensity in WT platelets stimulated with S100A8/A9 and H151. Data are presented as mean ± SD of 3-4 mice per group in at least two independent experiments. Statistical significance was determined using ordinary two-way ANOVA followed by Fisher’s least significant difference (LSD) multiple-comparisons test for experiments involving multiple groups and repeated-measures one-way ANOVA followed by Dunnett’s multiple-comparisons test for **e,f**. Comparisons between unstimulated and stimulated platelets from the same mouse were analyzed using paired statistical tests, as indicated in the figure. *P* < 0.05 (*), *P* < 0.01 (**); ns, not significant.

### TLR4 stimulation induces MCU-dependent mitochondrial depolarization and cytosolic mtDNA accumulation

Because LPS does not directly activate cGAS, we sought to identify the endogenous signal linking TLR4 stimulation to cGAS-STING activation. Prior studies have reported that TLR4 stimulation promotes mitochondrial depolarization and reactive oxygen species (ROS) production in platelets^43^. TLR signaling is also known to trigger intracellular calcium mobilization^44-48^ and subsequent mitochondrial calcium uptake through the MCU, an event previously reported to induce mitochondrial depolarization and ROS production in platelets^49-51^. Given that mitochondria represent a major source of immunostimulatory DNA that can be released upon calcium uptake by the mitochondria^52^, we investigated whether TLR4 stimulation induces MCU-dependent mitochondrial changes characterized by increased mitochondrial ROS (mtROS) production and membrane depolarization. Pharmacological inhibition of the MCU using RU360 during LPS stimulation significantly attenuated both responses (Fig. 5a, b, c). In addition, when we directly analysed platelets from WT mice post-LPS administration, we observed that mtROS were also increased *in vivo* (Fig. 5d), consistent with a change in mitochondrial functions as previously reported to occur during endotoxemia^6,8^. Consistent with mitochondria serving as a source of endogenous danger signals^21,22^, LPS also promoted accumulation of mtDNA in the cytosol, an effect that was markedly reduced by RU360 treatment (Fig. 5e). To directly assess whether MCU-dependent mitochondrial calcium uptake contributes to platelet activation downstream of TLR4 signaling, we measured CD62P surface expression in platelets stimulated with LPS alone or in the presence of RU360. Inhibition of the MCU significantly reduced LPS-induced CD62P expression (Fig. 5f), indicating that mitochondrial calcium uptake and mtDNA release are required for activation of platelets downstream TLR4 stimulation. Together, these findings identify MCU-dependent mitochondrial depolarization and mtDNA release as a mechanistic link between TLR4 stimulation and activation of the platelet cGAS-STING signaling pathway.

**Fig. 5.**
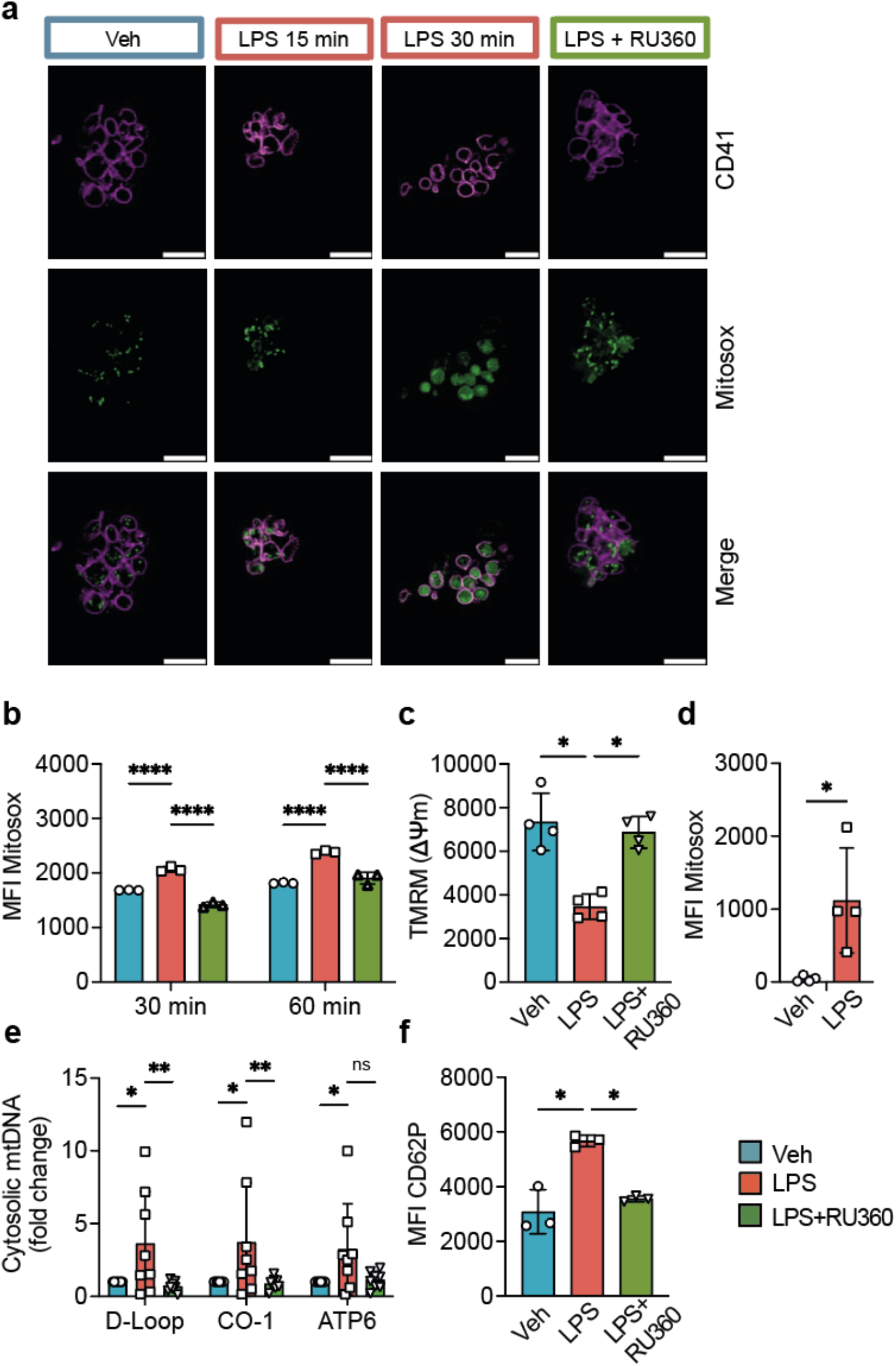
LPS induces MCU-dependent mitochondrial stress and cytosolic mtDNA accumulation in platelets. Washed WT platelets were stimulated with LPS in the presence or absence of the MCU inhibitor RU360, as indicated. **a**, Representative confocal images of mitochondrial ROS production following LPS stimulation for 15 or 30 min, or following co-treatment with LPS and RU360 for 30 min. **b**, Quantification of mtROS measured by flow cytometry following LPS stimulation for 30 min, or 1 hour. **c**, Mitochondrial membrane potential (ΔΨm) measured by TMRM fluorescence following stimulation with LPS in the presence or absence of RU360 for 1 hour. **d**, Mitochondrial ROS production in platelet from WT mice measured by flow cytometry 1 hour after intraperitoneal LPS injection (10 mg/kg). **e**, Relative abundance of cytosolic mitochondrial DNA measured by quantitative PCR in digitonin-isolated cytosolic fractions following stimulation with LPS in the presence or absence of RU360 for 1 hour. Mitochondrial DNA was quantified using primers targeting MT-*Atp6*, MT-*Co1*, and the MT-*D-loop* region. **f**, Platelet activation measured by CD62P expression following stimulation with LPS in the presence or absence of RU360 for 1 hour. Data are presented as mean ± SD. Each symbol represents one biologically independent mouse. *n* = 3-8 mice per group collected across at least two independent experiments. Statistical significance was determined using ordinary two-way ANOVA followed by Fisher’s least significant difference (LSD) (b, e); multiple-comparisons test for experiments involving multiple groups and repeated-measures one-way ANOVA followed by Dunnett’s multiple-comparisons test (c, f); and using an ordinary two-tailed unpaired Mann-Whitney test for (d). *P* < 0.05 (*)*, P < 0.01(**), P* < 0.0001 (****); ns, not significant. Image scale bar: 5 μm.

### The mitochondrial mtDNA-cGAS pathway extends beyond TLR4 stimulation

Given that TLR4 stimulation induced MCU-dependent mitochondrial depolarization, mtROS production and cytosolic mtDNA accumulation, we hypothesized that other innate immune receptors capable of mobilizing intracellular calcium might engage the same pathway. To test whether mitochondrial stress and mtDNA release represent a common signaling mechanism downstream of innate immune receptors, we examined platelet responses to TLR1/2 and TLR5 stimulation, since both cell membrane TLRs were reported to be present on platelets^4,11-13^.

The TLR1/2 agonist Pam3CSK4 induced robust platelet activation in WT platelets but failed to significantly activate *Cgas^-/-^* platelets, resulting in a significant increase in CD62P surface expression on WT platelets but not on *Cgas^-/-^* platelets (Fig. 6a). Similarly, stimulation with the TLR5 agonist flagellin increased platelet activation in WT platelets, although to a lesser extent than Pam3CSK4, and this response tended to be attenuated in *Cgas^-/-^* platelets (Fig. 6b). Together, these findings indicate that cGAS-dependent signaling is not restricted to TLR4 but also occurs downstream of other TLRs, suggesting convergence of distinct innate immune receptors on a shared signaling pathway.

**Fig. 6.**
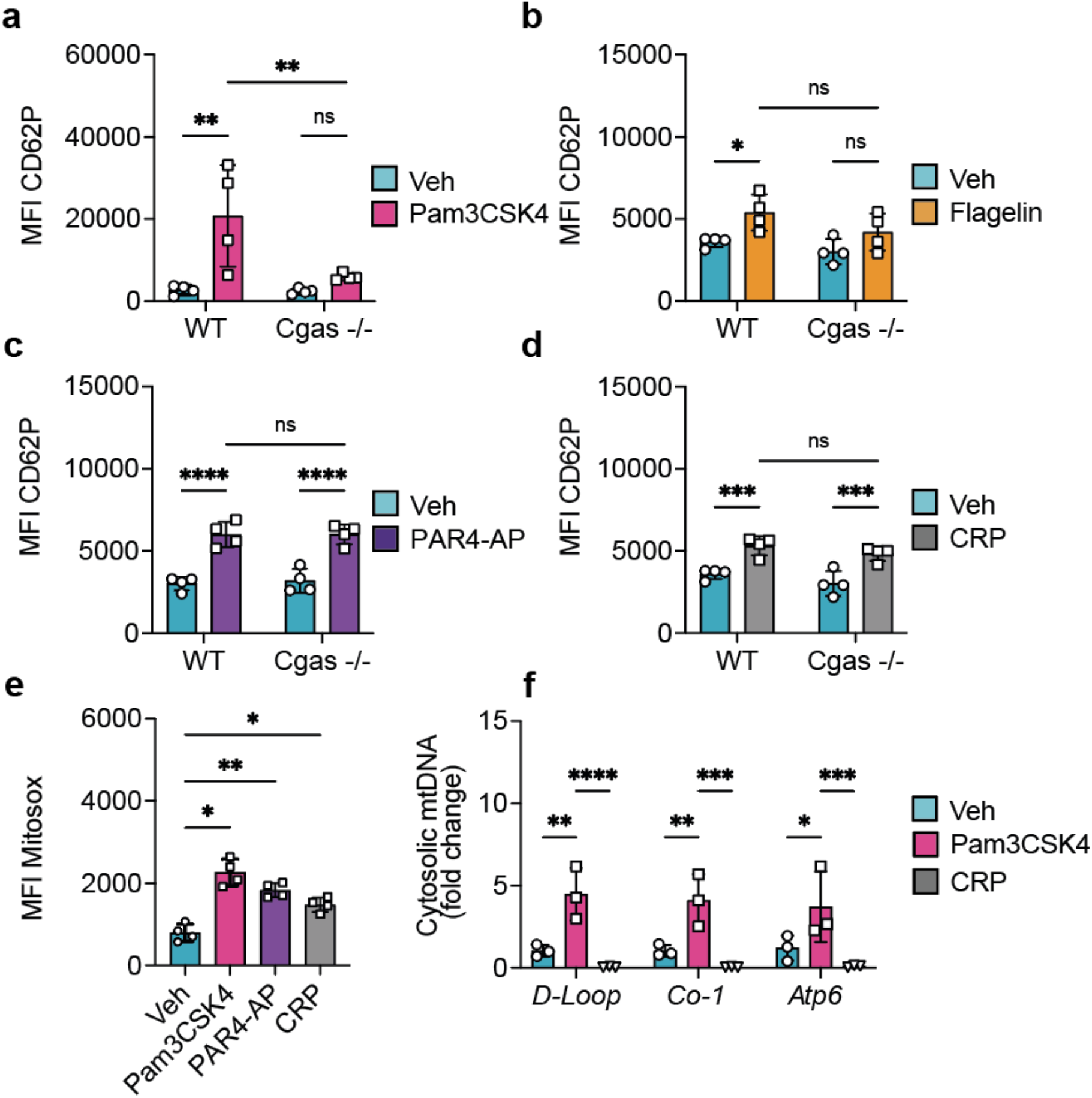
Multiple innate immune receptors converge on a common mitochondrial mtDNA-cGAS pathway. Washed WT and *Cgas*-/-platelets were stimulated as indicated and platelet activation, mitochondrial stress responses, and cytosolic mtDNA accumulation were assessed. **a**, Surface CD62P expression following stimulation with the TLR1/2 agonist Pam3CSK4 for 1 hour. **b**, Surface CD62P expression following stimulation with the TLR5 agonist flagellin for 1 hour. **c**, Surface CD62P expression following stimulation with PAR4-AP for 1 hour. **d**, Surface CD62P expression following stimulation with CRP for 1 hour. **e**, Mitochondrial ROS production measured by flow cytometry following stimulation with Pam3CSK4, PAR4-AP, or CRP for 1 hour. **f**, Relative abundance of cytosolic mtDNA measured by quantitative PCR in digitonin-isolated cytosolic fractions following stimulation with Pam3CSK4 or CRP for 1 hour. mtDNA was quantified using primers targeting MT-*Atp6*, MT-*Co1*, and the MT-D-loop region. Data are presented as mean ± SD. Each symbol represents one biologically independent mouse. *n* = 3-4 biologically independent mice per group collected across at least two independent experiments. Statistical significance was determined using ordinary two-way ANOVA followed by Fisher’s least significant difference (LSD) multiple-comparisons test for experiments involving multiple groups and repeated-measures one-way ANOVA followed by Dunnett’s multiple-comparisons test for **e**. *P* < 0.05 (\**), P < 0.01 (**), P < 0.001 (\*\*\**), *P* < 0.0001 (****); ns, not significant.

In contrast, platelet activation measured by P-selectin surface translocation, and induced by the canonical hemostatic agonists PAR4-activating peptide (PAR4-AP, AYPGKF-OH) and Collagen-Related Peptide (CRP) was fully preserved in *Cgas^-/-^* platelets (Fig. 6c, d), aligning with previous results showing that classical agonists do not directly engage cGAS-STING^24,25^. Thus, cGAS is dispensable for platelet activation by these canonical agonists but selectively contributes to platelet responses elicited by the innate immune receptor agonists examined here. To next investigate whether activation of distinct TLRs converged on a common mitochondrial pathway, we assessed mtROS and mtDNA release. Similar to LPS, Pam3CSK4 stimulation increased mtROS production and promoted accumulation of cytosolic mtDNA, as detected by ATP6, CO1 and D-loop sequences (Fig. 6e, f). In contrast, although CRP also induced mtROS production, CRP stimulation did not result in increased cytosolic mtDNA accumulation (Fig. 6e, f). These findings indicate that innate immune receptors, despite recognizing distinct ligands, converge on a common mitochondrial repolarization-mtDNA-cGAS signaling axis to promote platelet activation, whereas canonical hemostatic agonists trigger platelet activation through cGAS-independent mechanisms.

### Human platelets retain a selective innate immune mitochondrial-cGAS signaling axis

To determine whether this pathway is conserved in human platelets, we examined responses to innate immune and canonical platelet agonists in washed platelets isolated from healthy volunteers. Under the experimental conditions tested, LPS did not induce platelet aggregation, consistent with previous reports showing that LPS primarily potentiates platelet responses to classical agonists in human rather than directly activating platelets^53-55^. In contrast, Pam3CSK4 elicited robust and reproducible aggregation, leading us to focus our subsequent human platelet experiments on Pam3CSK4 (Fig. 7a). Pam3CSK4-induced CD62P expression was significantly reduced by pharmacological inhibition of STING with the small molecule inhibitor H151^42^ (Fig. 7b), indicating that cGAS-STING signaling contributes to TLR1/2-mediated platelet activation in human platelets. Pam3CSK4 also promoted mtROS production. Although this effect did not reach statistical significance, inhibition of the MCU with RU360 tended to attenuate the response (Fig. 7c). In contrast, stimulation with canonical platelet agonists PAR1-activating peptide (PAR1-AP; SFLLRN-NH2) and CRP induced platelet activation and mtROS production, but their effect on CD62P expression were only slightly affected by STING inhibition when compared with Pam3CSK4 (Fig. 7d, f), while mitochondrial ROS was not significantly reduced by MCU inhibition (Fig. 7e, g). In line with the involvement of STING in TLR1/2 mediated platelet activation, Pam3CSK4 stimulation also increased the abundance of cytosolic mtDNA accumulation, as indicated by elevated levels of the mitochondrial CO3 sequence. This response was prevented by MCU inhibition (Fig. 7h). Although the effect was less marked than in mouse platelets, stimulation with PAR1-AP or CRP did not result in a significant accumulation of cytosolic mtDNA, as measured by the mitochondrial CO3 quantification, compared with unstimulated controls (Fig. 7h). Together, these findings support conservation of a mtDNA-cGAS-STING signaling axis in human platelets following TLR1/2 stimulation and establish a mechanistic distinction between innate immune receptor-mediated and canonical platelet activation. Collectively, our data identify cGAS as a platelet-intrinsic sensor of mitochondria-derived DNA that functions as a signaling hub integrating innate immune receptor engagement with platelet activation and thromboinflammatory responses.

**Fig. 7.**
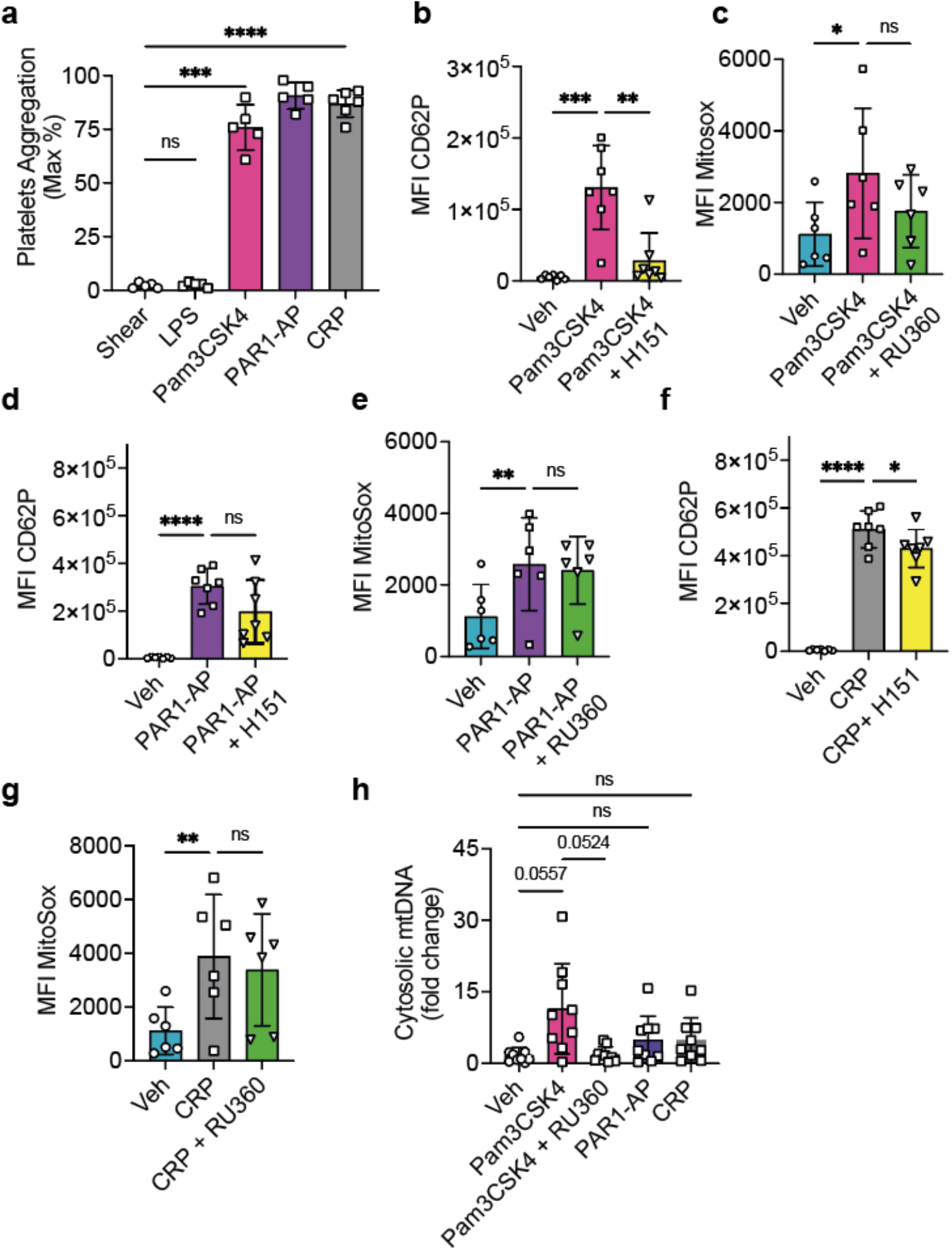
Human platelets retain an innate immune mitochondrial mtDNA-cGAS-STING signaling pathway. Washed human platelets were stimulated as indicated and platelet activation, aggregation, mitochondrial stress responses and cytosolic mtDNA accumulation were assessed. **a**, Maximal platelet aggregation measured by light transmission aggregometry following stimulation with LPS, Pam3CSK4, PAR1-AP, or CRP. **b**, Surface CD62P expression following stimulation with Pam3CSK4 in the presence or absence of the STING inhibitor H-151. **c**, Mitochondrial ROS production following stimulation with Pam3CSK4 in the presence or absence of the MCU inhibitor RU360. **d**, Surface CD62P expression following stimulation with PAR1-AP in the presence or absence of H-151. **e**, Mitochondrial ROS production following stimulation with PAR1-AP in the presence or absence of RU360. **f**, Surface CD62P expression following stimulation with CRP in the presence or absence of H-151. **g**, Mitochondrial ROS production following stimulation with CRP in the presence or absence of RU360. **h**, Relative abundance of cytosolic mtDNA measured by quantitative PCR in digitonin-isolated cytosolic fractions following stimulation with Pam3CSK4, PAR1-AP or CRP for 1 hour in the presence or absence of RU360. mtDNA was quantified using primers targeting MT-CO3. Data are presented as mean ± SD. Each symbol represents one biologically independent donor. *n* = 5-9 donors collected across at least five independent experiments. Statistical significance was determined using repeated-measures one-way ANOVA followed by Dunnett’s multiple-comparisons test. *P* < 0.05 (\**), P < 0.01 (**), P **<** 0.001 (\*\*\**), *P* < 0.0001 (****); ns, not significant.

## Discussion

Platelets contribute to host defense by sensing microbial and endogenous danger signals, yet the mechanisms that couple innate immune receptor engagement to platelet activation remain incompletely understood. Here, we identify a platelet-intrinsic mitochondrial stress-mtDNA-cGAS-STING signaling axis that connects innate immune receptor stimulation to platelet activation. We demonstrate that stimulation of TLR4, TLR1/2 and TLR5 induces MCU-dependent mitochondrial stress responses and cytosolic mtDNA accumulation, resulting in activation of cGAS-STING signaling. Functionally, this pathway amplifies platelet activation, inflammasome signaling and early inflammatory responses during endotoxemia. Together, our findings establish cGAS as a key component of inflammatory platelet signaling and reveal a mechanism through which distinct innate immune receptors converge to promote platelet activation.

The involvement of cGAS-STING signaling downstream of TLR stimulation contrasts with previous reports showing that direct pharmacological activation of this pathway is insufficient to induce platelet activation^24-26^. This suggests that cGAS-STING does not appear to function as an autonomous platelet activation pathway but rather cooperates with additional signaling mechanisms engaged downstream of TLRs. Although direct activation of cGAS-STING has been reported to potentiate platelet responses to canonical agonists^24,26^, TLR stimulation simultaneously engages receptor signaling pathways and promote mtDNA accumulation, thereby allowing cGAS-STING activation to amplify platelet activation. The molecular identity of the cooperating signals remains to be established but may involve calcium-dependent signaling, phosphoinositide 3-kinase (PI3K) pathways, or inflammasome-associated mechanisms activated downstream of TLR engagement^48,56,57^.

A central finding of this study is the identification of cGAS as a signal integration hub in platelets. In nucleated cells, cGAS is generally recognized as a cytosolic DNA sensor that activates innate immune signaling pathways, complementing TLR-mediated responses through the induction of type I interferons and inflammatory cytokines^58-60^. In contrast, our findings suggest a more central role in platelets, where multiple TLRs converge on cGAS-dependent signaling to promote activation. Rather than directly sensing microbial products, cGAS responded to mitochondria-derived DNA that accumulates in the cytosol following innate immune receptor stimulation. Despite recognizing distinct ligands, TLR4, TLR1/2 and TLR5 all engaged to a different extent a common mtDNA-cGAS sensing pathway, whereas canonical agonists including thrombin receptor-activating peptide (TRAP) and CRP activated platelets independently of cGAS and failed to induce substantial cytosolic mtDNA accumulation. Collectively, these observations suggest that cGAS does not function as a universal platelet activation mediator but rather constitutes a specialized signaling module that selectively couples innate immune engagement to inflammatory platelet responses.

While mtROS have been implicated as a key driver of mtDNA release in several cellular systems^61-63^, our findings suggest that mtROS production alone is insufficient to promote cytosolic mtDNA accumulation in platelets. Consistent with this notion, both TLR agonists and canonical agonists induced mtROS production, yet TLR stimulation predominantly resulted in substantial accumulation of cytosolic mtDNA. We therefore propose that mtDNA acts as a common intracellular danger signal through which distinct innate immune receptors converge on a shared platelet activation pathway. Notably, the magnitude of this response varied among TLR agonists, with Pam3CSK4 inducing a more pronounced cGAS-dependent activation than LPS or flagellin. These differences may reflect variations in receptor expression levels, signaling efficiency, or the extent of mitochondrial responses induced by individual TLR agonists. Differences in platelet TLR expression, which may also vary among individuals, could contribute to the heterogeneity of these responses^64,65^.

Our findings also extend the current understanding of platelet STING signaling. A previous study proposed that STING activation in platelets during sepsis primarily depends on the uptake of extracellular cGAMP produced by surrounding vascular and immune cells^24^. While this mechanism likely contributes to platelet activation *in vivo*, our data reveal that platelets can engage cGAS-dependent signaling autonomously, without requiring exogenous cGAMP or DNA administration^24,25^. Platelet mitochondrial depolarization and cGAS activation have also been reported in the context of hypoxia-associated periodontal inflammation^28^, but not as a consequence of direct signaling responses of innate immune surface receptors. Importantly, transfusion of WT, but not cGAS-deficient, platelets into thrombocytopenic *Cgas^-/-^* recipients, thus unable to produce cGAMP, restored platelet activation and amplified systemic inflammatory responses following endotoxin challenge, supporting a direct contribution of donor platelet-intrinsic cGAS signaling *in vivo*. Notably, the impact of cGAS deficiency was most pronounced during the early stages of endotoxemia and became less evident at later time points. This temporal pattern suggests that platelet-intrinsic cGAS initiates early inflammatory responses, whereas cGAMP produced by neighboring vascular or immune cells may increasingly contribute to STING activation as inflammation progresses^24,60^. Rather than representing mutually exclusive mechanisms, these findings support a model in which platelet STING integrates both intrinsic sensing of mitochondria-derived DNA and extrinsic signals originating from the surrounding inflammatory environment.

Mitochondrial membrane depolarization and mtROS production have emerged as hallmarks of platelet activation during sepsis and other inflammatory conditions^8,9,29,49,66^. Consistent with previous observations, innate immune receptor stimulation induced mtROS production, mitochondrial membrane depolarization and accumulation of cytosolic mtDNA. Importantly, pharmacological inhibition of the MCU attenuated these mitochondrial responses and reduced mtDNA release, placing mitochondrial calcium uptake upstream of mtDNA-cGAS signaling. Interestingly, the MCU has previously been implicated in platelet activation and metabolism, including signaling downstream of the immunoreceptor tyrosine-based activation motif (ITAM)-associated receptors GPVI and CLEC-2^49-51^. However, despite evidence that MCU contributes to GPVI-mediated signaling, CRP stimulation did not induce significant cytosolic mtDNA accumulation *in vitro*. These results suggest that mitochondrial calcium uptake is necessary but not sufficient for engagement of the mtDNA-cGAS pathway and that innate immune receptors integrate signaling specificities to promote mitochondrial danger signaling.

The molecular mechanisms responsible for cytosolic mtDNA release from mitochondria remain incompletely understood. Several pathways, including mitochondrial permeability transition pore (mPTP) opening, voltage-dependent anion channel (VDAC) oligomerization and BAX/BAK-mediated mitochondrial membrane permeabilization, have been implicated in mtDNA release in other cellular systems^9,62,63,67^. Because MCU inhibition reduced cytosolic mtDNA accumulation in our study, calcium-mediated mitochondrial permeability mechanisms may represent a particularly plausible mechanism linking TLR stimulation and mtDNA-dependent cGAS signaling in platelets.

While the endotoxemia model used in this study recapitulates several key inflammatory features of sepsis and our findings provide important mechanistic insights into platelet innate immune signaling, several limitations should be acknowledged. First, depletion of endogenous platelets using anti-platelet antibodies, which was required to perform platelet transfer experiments, precluded longitudinal assessment of platelet function *in vivo* and limited our ability to evaluate the temporal contribution of platelet cGAS throughout the course of endotoxemia. Second, we did not correlate platelet responses in human donors with the expression levels of TLRs, which may have contributed to the inter-individual variability observed in some experiments. Finally, although our data place MCU-dependent mitochondrial responses upstream of cGAS-dependent platelet activation, the additional signaling events that couple TLR engagement to mtDNA release remain incompletely defined and warrant further investigation.

In summary, we identify a platelet-intrinsic mitochondrial stress-mtDNA-cGAS-STING signaling axis that links innate immune receptor stimulation to platelet activation. By sensing endogenous mtDNA accumulation following mitochondrial stress, cGAS emerges as an intracellular integrator of inflammatory signaling in platelets. These findings reveal a previously unrecognized mechanism through which distinct innate immune receptors converge to amplify platelet responses and establish the mtDNA-cGAS axis as a key contributor to platelet-mediated thromboinflammation.

## METHODS

### Animals

Mice were maintained under specific pathogen-free conditions with *ad libitum* access to food and water at the animal facility of the Université du Québec à Trois-Rivières. Mice were housed under a 12-h reversed light-dark cycle with lights off at 07:00 and lights on at 19:00. Wild-type (WT) C57BL/6J mice and *Cgas^-/-^*mice (B6(C)-Cgastm1d(EUCOMM)Hmgu/J) were obtained from The Jackson Laboratory. Mice aged 8-14 weeks were used for *in vitro* studies. Male mice aged 8-10 weeks were used for endotoxemia experiments. Genotypes were confirmed by PCR and loss of cGAS expression in platelets was verified by western blot analysis as previously described^26^. All animal procedures were approved by the institutional animal care committee of the Université du Québec à Trois-Rivières (protocols 2021-G.P.1, 2023-G.P.11 and 2026-G.P.15) and were conducted in accordance with Canadian Council on Animal Care guidelines.

### Endotoxemia model

Experimental endotoxemia was induced by intraperitoneal administration of lipopolysaccharide (LPS E. Coli 0111:B4, Sigma, L2630) at 10 mg/kg in 150µl. Control animals received an equivalent volume of sterile PBS (vehicle). Blood was collected at the indicated time points for assessment of platelet activation, platelet-neutrophil aggregate formation, cytokine production and platelet counts.

### Platelet depletion and adoptive platelet transfer

Platelet depletion was performed using a rabbit anti-mouse thrombocyte polyclonal antiserum, based on previously described antibody-mediated depletion approach^41^. Briefly, recipient mice received 7.5 μL anti-mouse thrombocyte antiserum (Cedarlane, CLA31440) diluted with PBS to a final volume of 150 μL by intraperitoneal injection. Platelet counts were assessed 12 hours after antiserum administration to confirm depletion. At this time point, circulating platelet counts were reduced to 6.8 ± 9.8% of baseline levels. Following platelet depletion, *Cgas^-/-^* recipient mice received 100 µL of isolated platelets from either WT or *Cgas^-/-^* donor mice via tail vein injection. Donor platelets were at a concentration of 5 × 10^8^ platelets/mL corresponding to a fixed dose of 5 × 10^7^ platelets per recipient mouse. Platelet counts were determined in whole blood using a CytoFLEX flow cytometer (Beckman Coulter). One hour after platelet transfusion, recipient mice were challenged with LPS (10 mg/kg) and analyzed at the indicated time points.

### Mouse platelets isolation

Whole blood (0.8-1ml per mouse) was collected into acid-citrate-dextrose (ACD) anticoagulant at a blood-to-ACD ratio of 10:1 (v/v). Platelet-rich plasma (PRP) was obtained by centrifugation at 300 x g for 8 min. Residual leukocytes and erythrocytes were removed by a second centrifugation step at 150 × g for 4 min. PRP was subsequently centrifuged at 800 x g for 8 min to pellet platelets. Plasma was collected and stored at -80°C for subsequent analyses. Platelets were resuspended in Hank’s balanced salt solution (HBSS) at a concentration of 2.5 x 10^8^ platelets/mL. The platelet suspension was subsequently divided into separate aliquots for the indicated experimental conditions and immediately processed for *in vitro* stimulation or flow cytometric analysis and allowed to rest for 10 min at room temperature before stimulation. No PGE1 was used during mouse platelet isolation.

### Human platelets isolation

Venous blood was obtained from healthy adult volunteers following written informed consent. Donors were aged between 18 and 50 years of age, reported no regular medication use and had not taken aspirin-containing medication or drugs affecting platelet function for at least 3 weeks before blood collection, had not undergone major surgery in the previous 6 months, did not have a history of bleeding symptoms, and had platelet counts and hemoglobin levels within the normal reference range. Both male and female donors were included. All procedures involving human participants were approved by the Research Ethics Committee of Montreal Heart Institute (protocol 2017-2154), and were conducted in accordance with the Declaration of Helsinki. Blood was drawn by venipuncture into syringes containing acid citrate dextrose (ACD-A, DIN: 00788139) in a 1:5 volume ratio (ACD/blood). Blood was transferred to 50 mL tubes, centrifuged at 200 X g for 10 min without a break, and PRP was collected. Prostaglandin E1 (1 µM) was added to PRP prior to centrifugation at 1000 x g for 10 min. Plasma was removed and discarded. Pelleted platelets were resuspended in Tyrode’s buffer (137 mM NaCl, 11.9 mM NaHCO3, 0.4 mM NaH2PO4, 2.7 mM KCl, 1.1 mM MgCl2, 5.6 mM glucose, pH 7.4). Platelet counts were determined using a Beckman Coulter hematology analyzer (DxH 520) and adjusted to a final concentration of 2.5 x 10^8^/mL. Platelets were allowed to rest at room temperature for 60 min prior to functional experiments.

### Platelet stimulation

Washed platelets from mice were stimulated with LPS (10 μg/mL), recombinant S100A8/A9 heterodimer (10 μg/mL), Pam3CSK4 (10 μg/mL), flagellin (10 μg/mL), collagen-related peptide (CRP; 1 μg/mL) or thrombin receptor activating peptide for the indicated times, as appropriate for each experiment. Mouse platelets were stimulated with PAR4-activating peptide AYPGKF-OH (125 μM), and human platelets with PAR1-activating peptide SFLLRN-NH2 (3 μM). Where indicated, platelets were pre-incubated with the STING inhibitor H-151 (10 μM), the mitochondrial calcium uniporter inhibitor RU360 (10 μM), or vehicle control for 1 hour before stimulation. We used, LPS from Escherichia coli O111:B4 (Sigma-Aldrich, L2630), Pam3CSK4 (InvivoGen, tlrl-pms), flagellin (InvivoGen, tlrl-pbsfla), recombinant mouse S100A8/A9 heterodimer (R&D Systems, 8916-S8), cross-linked collagen-related peptide (CRP-XL; CambCol Laboratories Ltd., AB110421M), mouse PAR4-activating peptide AYPGKF-OH (Cayman Chemical, 27132), human PAR1-activating peptide TRAP-6 amide trifluoroacetate (SFLLRN-NH2; Bachem Americas, H-2936.0005BA) H-151 (MedChemExpress, HY-112693), and RU360 (MilliporeSigma, 557440) were used as indicated. In most experiment following stimulation, mouse platelets were fixed with 2% of paraformaldehyde for 15 min and washed before antibody staining prior flow cytometric analysis. For human platelet, a 5 µL of the stimulated platelet suspension was collected and stained directly without fixation.

### Flow cytometric analysis of platelet activation/aggregation and phosphatidylserine exposure

Platelet activation was assessed by measuring P-selectin (CD62P) surface expression by flow cytometry. Following initial gating based on forward-scatter and side-scatter characteristics, platelets were identified as CD41 positive events using Pacific Blue™ anti-mouse CD41 (Biolegend, clone MWReg30, 133932). Platelets were stained PE anti-mouse/rat CD62P (Biolegend, clone RMP-1, 148306) for 30 min at room temperature and CD62P expression was quantified as mean fluorescence intensity (MFI) within CD41 positive population. Platelet-neutrophil aggregates were identified using a sequential gating strategy. Following initial gating based on forward-scatter and side-scatter characteristics, CD41 positive events were selected, followed by sequential gating of CD11b positive and Ly6G positive events. Platelet aggregation with neutrophils was quantified as CD41 MFI within the Ly6G positive population and platelet aggregation with monocytes within Ly6G negative population. Cleaved Caspase-1 was measured using the Geen Pyroptosis/Caspase-1 Assay (ImmunoChemistry Technologies, 9146) according to the manufacturer’s instructions, and fluorescence was quantified within the platelet CD41 positive population. Phosphatidylserine exposure was determined using either the Annexin V-FITC Apoptosis Detection Kit (Invitrogen; BMS500FI-20) or the Annexin V-FITC Apoptosis detection Kit (Cayman Chemical, 600300), according to the manufacturers’ instructions. Annexin V-FITC fluorescence was quantified as MFI within the CD41 positive platelet population. Human reagents for flow cytometry were: CD62P VioBright V515 (Miltenyi Biotec, 130-130-572), CD41/CD61 APC Vio770 (Miltenyi Biotec, 130-119-574), CD41/CD61 FITC (Miltenyi Biotec, 139-124-887) and Annexin V Alexa Fluor 647 (Invitrogen, A23204A). A minimum of 5000 events were acquired per sample using a CytoFLEX flow cytometer for mouse platelets (Beckman Coulter) and a spectral flow cytometer Northern light (Cytek Biosciences) for human platelets. Data were analyzed using FlowJo software (version 10.10.1).

### Light transmission aggregometry

Human platelet aggregation was assessed by light transmission aggregometry using a Chrono-Log Model 700 aggregometer (Chrono-Log, USA) at 37°C under continuous stirring (1,200 rpm). Aggregation traces were recorded using AGGRO/LINK software. Platelets were stimulated with cross-linked collagen-related peptide (CRP-XL 1 μg/mL), PAR1-AP TRAP-6 (SFLLRN-NH2; 3 μM), Pam3CSK4 (10 μg/mL) or LPS (10 μg/mL). Where indicated, platelets were pre-incubated with H-151 (10 μM), RU360 (10 μM), or corresponding vehicle control for 1 hour before stimulation.

### Cytokine and plasma mediator quantification

Plasma concentrations of IL-1β (R&D, DY401), IL-6 (BD Biosciences, 555240), TNF (Abcam, 430904), CCL5 (R&D, DY478), S100A8/A9 (R&D, DY8596) and thrombin-antithrombin complexes (TAT) (Abcam, ab137994) were quantified using commercially available ELISA kits according to the manufacturers’ instructions.

### mtDNA quantification in platelet

Platelet cytosolic fractions were isolated using digitonin-mediated plasma membrane permeabilization as previously described with minor modifications^68,69^. Platelets isolated from two mice were pooled for each biological replicate and were separated into three tubes, two of which were used for cytosolic fractionation and one as a total mtDNA control. Platelets were stimulated as described above for 60 minutes before cytosolic fractionation. Briefly, platelets were incubated in 500 µL of buffer containing 30 µg/mL digitonin (Sigma-Aldrich, D141) for 10 minutes at 37 °C with gentle agitation at 300 rpm. Samples were centrifuged at 2,500 x g for 8 minutes at RT, after which 250 μL of the supernatant was transferred to a new tube and centrifuged at 10,000 x g for 8 minutes at RT. Subsequently, 100 µL of the resulting supernatant was transferred to a new tube and centrifuged again at 10,000 x g for 8 minutes at RT. A 25 µL aliquot of the final supernatant was collected as the cytosolic fraction and stored at -80 °C until quantitative PCR analysis. The total platelet DNA was extracted from the corresponding unfractionated platelet tube and used as the reference for relative mtDNA quantification. Relative mtDNA abundance was quantified by qPCR using primers targeting MT-CO3, MT-*Co1*, MT-*D-loop* and MT-*Atp6* on a CFX Real-Time PCR Detection System (Bio-Rad).

Human MT-CO3

Forward: ATGACCCACCAATCACATGC

Reverse: ATCACATGGCTAGGCCGGAG

Mouse MT-*Co1*

Forward: GCCCCAGATATAGCATTCCC

Reverse: GTTCATCCTGTTCCTGCTCC

Mouse MT-*D-loop*

Forward: AATCTACCATCCTCCGTGAAACC

Reverse: TCAGTTTAGCTACCCCCAAGTTTAA

Mouse MT-*Atp6*

Forward: TTCCCATCCTCAAAACGCCTAA

Reverse: GAAGTGGGCAAGTGAGCTTTT

Relative mtDNA abundance was quantified using the 2^-ΔΔCt^ method^70^. Cytosolic mtDNA signals were normalized to total platelet DNA and expressed relative to unstimulated controls.

### Measurement of mitochondrial reactive oxygen species

Mouse platelets were incubated with 500 nM MitoSOX Red superoxide indicator (Invitrogen, M36007) for 30 min at 37°C. For the 15 min condition, platelets were first incubated with MitoSox for 15 min, then stimulated with LPS (10 µg/mL) for an additional 15 min in the presence of MitoSox. For the 30 min condition, MitoSox and LPS were incubated simultaneously for 30 min. For the 60 min condition, platelets were stimulated with LPS for 30 min before adding MitoSox for the remaining 30 min stimulation. Following stimulation, platelets were fixed with 2% paraformaldehyde for 15 min, washed and stained with Pacific Blue anti-CD41 antibody before analysis by flow cytometry. The same protocol was used for stimulation with CRP-XL (1µg/mL), PAR4-AP (125 µM), or Pam3CSK4 (10 µg/mL) with or without 10 µM MCU inhibitor RU360 before stimulation. Washed human platelets were stimulated with indicated agonists and subsequently incubated with 500 nM MitoSox at 37 °C for 30 min before flow cytometry analyses. Where indicated, platelets were pre-treated with RU360 (10 μM) before stimulation.

### Confocal microscopy

For confocal imaging of mtROS, platelets were stained with MitoSox as described above and identified with CD41-Alexa Fluor 647. Coverslips were sealed using VALAP. Images were acquired as Z-stacks using a STELLARIS 5 confocal microscope model equipped with a DMi8 inverted microscope platform and a x60 objective (Leica Microsystems CMS GmbH). Images acquisitions and analyses were performed using LAS X and ImageJ software.

### Measurement of mitochondrial membrane potential

Mitochondrial membrane potential (ΔΨm) was measured using tetramethyl rhodamine methyl ester perchlorate (TMRM; Invitrogen, T668). Mouse platelets were incubated with TMRM (500 nM) and LPS (10 μg/mL) simultaneously for 60 min at 37°C. Following stimulation platelets were fixed with paraformaldehyde for 15 min, washed, stained with Pacific Blue anti-CD41 antibody, and analyzed by flow cytometry. Where indicated, platelets were pre-incubated with RU360 (10 μM) before stimulation. TMRM MFI was quantified within CD41 positive platelets population as a measure of ΔΨm.

### Statistical analysis

Statistical analyses were performed using GraphPad Prism version 11 (GraphPad Software). Data are presented as mean ± standard deviation (SD). Statistical tests used for each experiment are specified in the corresponding figure legends. Two-tailed tests were used throughout and P < 0.05 after correction for multiplicity was considered statistically significant. Exact sample sizes (n) are reported in the figure legends. For murine *in vivo* and *in vitro* experiments, or experiments with human platelets, n refer to individual mice, independent biological preparations, or independent healthy volunteers respectively. Technical replicates were not considered independent biological replicates.

## ACKNOWLEDGEMENT

We thank Melodie B. Plourde for training AB and LA on the instruments; Nadia Desnoyers, Michel Demers and Maude Whitton for help and support with the animal work.

## AUTHOR CONTRIBUTIONS

AB performed the experiments, analyzed data and wrote the first draft of the manuscript. LA, SF, MW, MO performed some experiments, analyzed data and reviewed the manuscript. EP and MG advise on research plan and edited the manuscript. ML supervised the work with human platelets and reviewed the manuscript. GP conceived and coordinated the study, designed and analyzed experiments, and finalized the manuscript. All authors reviewed the results and approved the final version of the manuscript.

---

**Supplementary Fig. 1.**
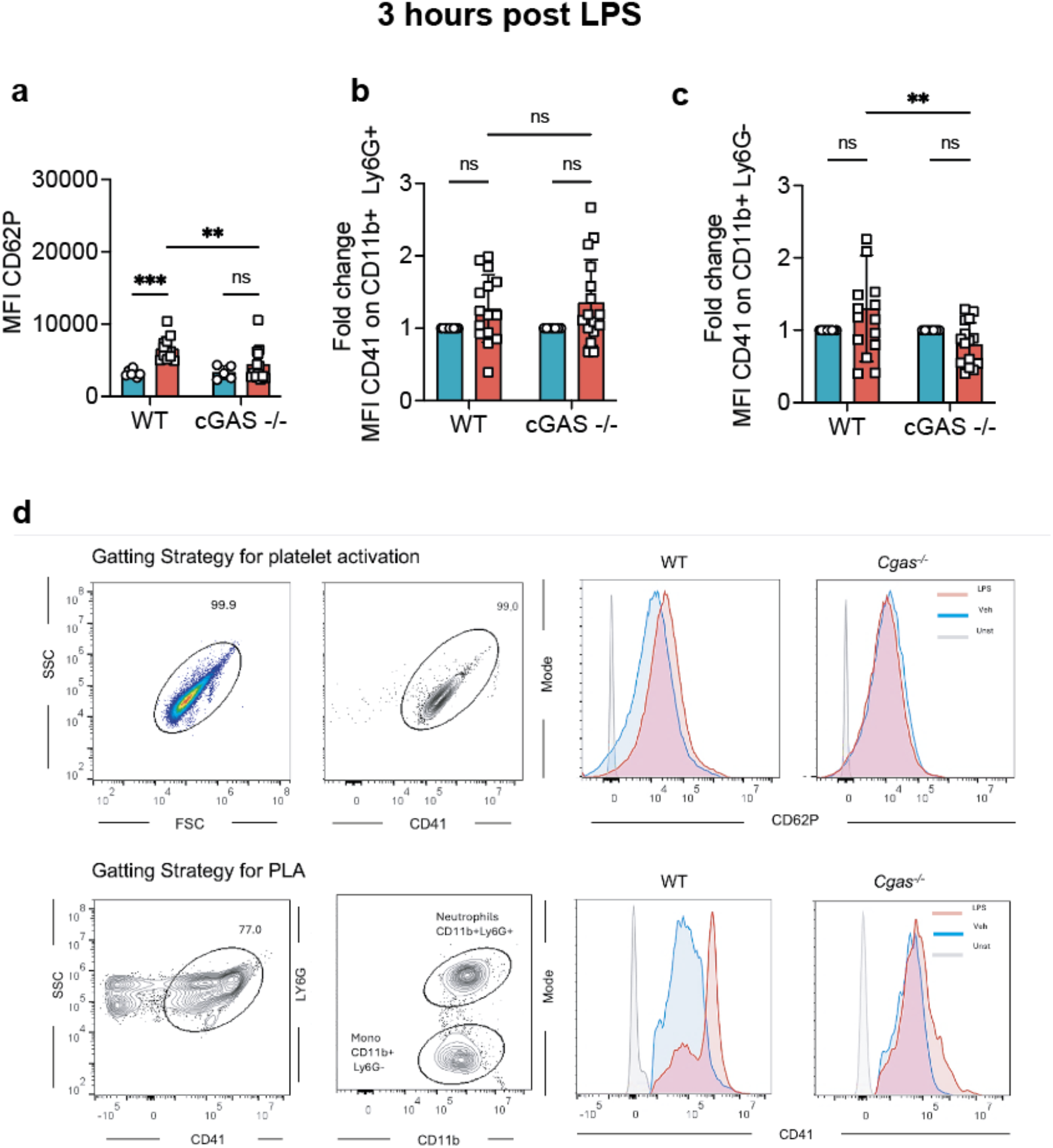
**a-d**, Platelet responses measured 3 hours following intraperitoneal injection of LPS (10 mg/kg) or saline in WT and *Cgas*^-/-^ mice. **a**, Platelet activation quantified by surface CD62P expression on circulating CD41 platelets population. **b**, Platelet-neutrophil aggregates quantified as CD41 fluorescence intensity on CD11b positive Ly6G positive population. **c**, Platelet-monocyte aggregates quantified as CD41 fluorescence intensity on CD11b positive Ly6G negative population. d. Gating strategy to measure platelet activation and PLA. Data are presented as mean ± SD. Each symbol represents one mouse. *n* = 5-16 biologically independent mice per group. Statistical significance was determined using ordinary two-way ANOVA followed by Fisher’s least significant difference (LSD) multiple-comparisons test. **P < 0.01; ***P < 0.001; ns, not significant.

**Supplementary Fig. 2.**
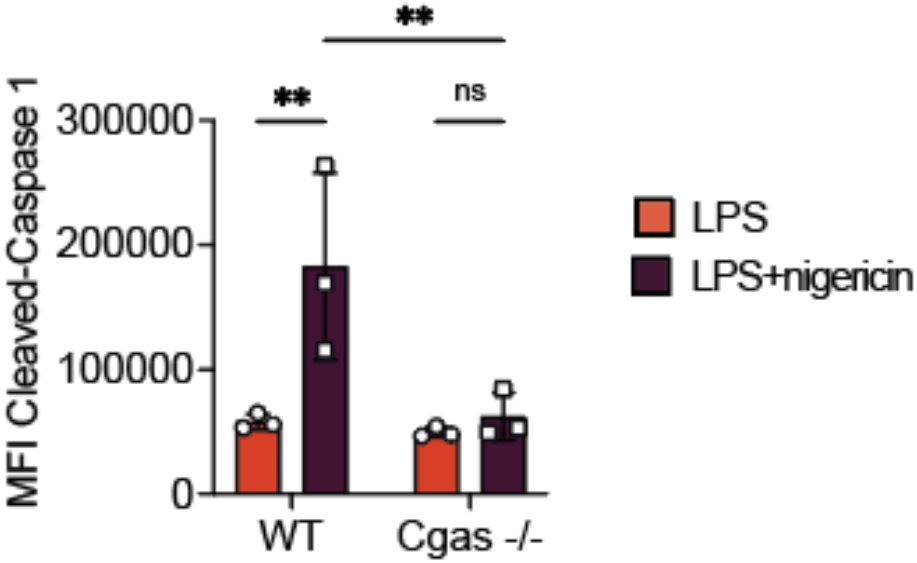
Washed mouse platelets were stimulated *in vitro* as indicated. Cleaved Caspase-1 measured by FLICA mean fluorescence intensity in WT and *Cgas*^-/-^ platelets following stimulation with LPS alone or with nigericin. Data are presented as mean ± SD of 3 mice per group in at least two independent experiments. Statistical significance was determined using ordinary two-way ANOVA followed by Fisher’s least significant difference (LSD) multiple-comparisons test. *P* < 0.01 (**); ns, not significant.

## References

1 Nicolai, L., Pekayvaz, K. & Massberg, S. Platelets: Orchestrators of immunity in host defense and beyond. Immunity 57, 957–972, doi:10.1016/j.immuni.2024.04.008 (2024).

2 Escaig, R., Kreutz, L., Kaiser, R. & Nicolai, L. Platelet effector functions in inflammation. Res Pract Thromb Haemost 10, 103412, doi:10.1016/j.rpth.2026.103412 (2026).

3 Cognasse, F. et al. The Inflammatory Role of Platelets via Their TLRs and Siglec Receptors. Front Immunol 6, 83, doi:10.3389/fimmu.2015.00083 (2015).

4 Blair, P. et al. Stimulation of Toll-like receptor 2 in human platelets induces a thromboinflammatory response through activation of phosphoinositide 3-kinase. Circ Res 104, 346–354, doi:10.1161/CIRCRESAHA.108.185785 (2009).

5 Marin Oyarzun, C. P., et al. Platelet Toll-Like Receptors Mediate Thromboinflammatory Responses in Patients With Essential Thrombocythemia. Front Immunol 11, 705, doi:10.3389/fimmu.2020.00705 (2020).

6 Su, M. et al. Gasdermin D-dependent platelet pyroptosis exacerbates NET formation and inflammation in severe sepsis. Nat Cardiovasc Res 1, 732–747, doi:10.1038/s44161-022-00108-7 (2022).

7 Clark, S. R. et al. Platelet TLR4 activates neutrophil extracellular traps to ensnare bacteria in septic blood. Nat Med 13, 463–469, doi:10.1038/nm1565 (2007).

8 Li, Y. & Feng, G. TLR4 inhibitor alleviates sepsis-induced organ failure by inhibiting platelet mtROS production, autophagy, and GPIIb/IIIa expression. J Bioenerg Biomembr 54, 155–162, doi:10.1007/s10863-022-09940-9 (2022).

9 Chen, C. C. et al. Lipopolysaccharide Potentiates Platelet Aggregation in Association with Apoptosis Through a Novel TLR4-Bax/Bcl-2-Mitochondrial Dysfunction Axis in Humans. Biomolecules 15, doi:10.3390/biom15121638 (2025).

10 Claushuis, T. A. M. et al. Platelet Toll-like receptor expression and activation induced by lipopolysaccharide and sepsis. Platelets 30, 296–304, doi:10.1080/09537104.2018.1445841 (2019).

11 Keane, C. et al. Invasive Streptococcus pneumoniae trigger platelet activation via Toll-like receptor 2. J Thromb Haemost 8, 2757–2765, doi:10.1111/j.1538-7836.2010.04093.x (2010).

12 Liu, X. et al. Strains of Group B streptococci from septic patients induce platelet activation via Toll-like Receptor 2. Clin Exp Pharmacol Physiol 44, 335–343, doi:10.1111/1440-1681.12707 (2017).

13 Williams, B. et al. Toll-like receptors 2 and 7 mediate coagulation activation and coagulopathy in murine sepsis. J Thromb Haemost 17, 1683–1693, doi:10.1111/jth.14543 (2019).

14 Takeda, K. & Akira, S. TLR signaling pathways. Semin Immunol 16, 3–9, doi:10.1016/j.smim.2003.10.003 (2004).

15 Fitzgerald, K. A. & Kagan, J. C. Toll-like Receptors and the Control of Immunity. Cell 180, 1044–1066, doi:10.1016/j.cell.2020.02.041 (2020).

16 Zhang, Z. & Zhang, C. Regulation of cGAS-STING signalling and its diversity of cellular outcomes. Nat Rev Immunol 25, 425–444, doi:10.1038/s41577-024-01112-7 (2025).

17 Zhang, B., Xu, P. & Ablasser, A. Regulation of the cGAS-STING Pathway. Annu Rev Immunol 43, 667–692, doi:10.1146/annurev-immunol-101721-032910 (2025).

18 Sun, L., Wu, J., Du, F., Chen, X. & Chen, Z. J. Cyclic GMP-AMP synthase is a cytosolic DNA sensor that activates the type I interferon pathway. Science 339, 786–791, doi:10.1126/science.1232458 (2013).

19 Zhang, X. et al. Cyclic GMP-AMP containing mixed phosphodiester linkages is an endogenous high-affinity ligand for STING. Mol Cell 51, 226–235, doi:10.1016/j.molcel.2013.05.022 (2013).

20 Ablasser, A. et al. cGAS produces a 2’-5’-linked cyclic dinucleotide second messenger that activates STING. Nature 498, 380–384, doi:10.1038/nature12306 (2013).

21 Hu, M. M. & Shu, H. B. Mitochondrial DNA-triggered innate immune response: mechanisms and diseases. Cell Mol Immunol 20, 1403–1412, doi:10.1038/s41423-023-01086-x (2023).

22 Giordano, L., Ware, S. A., Lagranha, C. J. & Kaufman, B. A. Mitochondrial DNA signals driving immune responses: Why, How, Where? Cell Commun Signal 23, 192, doi:10.1186/s12964-025-02042-0 (2025).

23 Harapas, C. R. et al. Organellar homeostasis and innate immune sensing. Nat Rev Immunol 22, 535–549, doi:10.1038/s41577-022-00682-8 (2022).

24 Yang, M. et al. STING activation in platelets aggravates septic thrombosis by enhancing platelet activation and granule secretion. Immunity 56, 1013–1026 e1016, doi:10.1016/j.immuni.2023.02.015 (2023).

25 Zhang, W. et al. Double-stranded DNA enhances platelet activation, thrombosis, and myocardial injury via cyclic GMP-AMP synthase. Cardiovasc Res 121, 353–366, doi:10.1093/cvr/cvae218 (2025).

26 El-Mortada, F. et al. Megakaryocytes possess a STING pathway that is transferred to platelets to potentiate activation. Life Sci Alliance 7, doi:10.26508/lsa.202302211 (2024).

27 Franco, C. V., Araya-Maturana, R. & Fuentes, E. Mitochondrial metabolism as a determinant of heterogeneous platelet functional phenotypes: Implications for antiplatelet resistance. Free Radic Biol Med 251, 466–479, doi:10.1016/j.freeradbiomed.2026.04.145 (2026).

28 Zhu, Y. et al. Platelet methyltransferase-like protein 4-mediated mitochondrial DNA metabolic disorder exacerbates oral mucosal immunopathology in hypoxia. Int J Oral Sci 17, 49, doi:10.1038/s41368-025-00373-9 (2025).

29 Ding, Y. et al. MTH1 protects platelet mitochondria from oxidative damage and regulates platelet function and thrombosis. Nat Commun 14, 4829, doi:10.1038/s41467-023-40600-7 (2023).

30 Danese, S. et al. Cutting edge: T cells trigger CD40-dependent platelet activation and granular RANTES release: a novel pathway for immune response amplification. J Immunol 172, 2011–2015, doi:10.4049/jimmunol.172.4.2011 (2004).

31 Klinger, M. H. et al. Immunocytochemical localization of the chemokines RANTES and MIP-1 alpha within human platelets and their release during storage. Int Arch Allergy Immunol 107, 541–546, doi:10.1159/000237097 (1995).

32 Pennings, G. J. et al. Rapid Release of Interleukin-1beta from Human Platelets Is Independent of NLRP3 and Caspase. Thromb Haemost 122, 517–528, doi:10.1055/s-0041-1731288 (2022).

33 Cha, J. K. et al. Activated platelets induce secretion of interleukin-1beta, monocyte chemotactic protein-1, and macrophage inflammatory protein-1alpha and surface expression of intercellular adhesion molecule-1 on cultured endothelial cells. J Korean Med Sci 15, 273–278, doi:10.3346/jkms.2000.15.3.273 (2000).

34 Rolfes, V. et al. Platelets Fuel the Inflammasome Activation of Innate Immune Cells. Cell Rep 31, 107615, doi:10.1016/j.celrep.2020.107615 (2020).

35 Yin, H. et al. Role for platelet glycoprotein Ib-IX and effects of its inhibition in endotoxemia-induced thrombosis, thrombocytopenia, and mortality. Arterioscler Thromb Vasc Biol 33, 2529–2537, doi:10.1161/ATVBAHA.113.302339 (2013).

36 Steven, S. et al. Time Response of Oxidative/Nitrosative Stress and Inflammation in LPS-Induced Endotoxaemia-A Comparative Study of Mice and Rats. Int J Mol Sci 18, doi:10.3390/ijms18102176 (2017).

37 Murthy, P. et al. The NLRP3 inflammasome and bruton’s tyrosine kinase in platelets co-regulate platelet activation, aggregation, and in vitro thrombus formation. Biochem Biophys Res Commun 483, 230–236, doi:10.1016/j.bbrc.2016.12.161 (2017).

38 Brown, G. T., Narayanan, P., Li, W., Silverstein, R. L. & McIntyre, T. M. Lipopolysaccharide stimulates platelets through an IL-1beta autocrine loop. J Immunol 191, 5196–5203, doi:10.4049/jimmunol.1300354 (2013).

39 Mason, K. D. et al. Programmed anuclear cell death delimits platelet life span. Cell 128, 1173–1186, doi:10.1016/j.cell.2007.01.037 (2007).

40 White, M. J. et al. Apoptotic caspases suppress mtDNA-induced STING-mediated type I IFN production. Cell 159, 1549–1562, doi:10.1016/j.cell.2014.11.036 (2014).

41 Salzmann, M. et al. Genetic platelet depletion is superior in platelet transfusion compared to current models. Haematologica 105, 1738–1749, doi:10.3324/haematol.2019.222448 (2020).

42 Haag, S. M. et al. Targeting STING with covalent small-molecule inhibitors. Nature 559, 269–273, doi:10.1038/s41586-018-0287-8 (2018).

43 Ip, W. K. E., Hoshi, N., Shouval, D. S., Snapper, S. & Medzhitov, R. Anti-inflammatory effect of IL-10 mediated by metabolic reprogramming of macrophages. Science 356, 513–519, doi:10.1126/science.aal3535 (2017).

44 Park, K. S. et al. TLR3-/4-Priming Differentially Promotes Ca(2+) Signaling and Cytokine Expression and Ca(2+)-Dependently Augments Cytokine Release in hMSCs. Sci Rep 6, 23103, doi:10.1038/srep23103 (2016).

45 Birla, H. et al. Toll-like receptor 4 activation enhances Orai1-mediated calcium signal promoting cytokine production in spinal astrocytes. Cell Calcium 105, 102619, doi:10.1016/j.ceca.2022.102619 (2022).

46 Ren, H. et al. Toll-like receptor-triggered calcium mobilization protects mice against bacterial infection through extracellular ATP release. Infect Immun 82, 5076–5085, doi:10.1128/IAI.02546-14 (2014).

47 Geng, J. et al. TLR4 signalling via Piezo1 engages and enhances the macrophage mediated host response during bacterial infection. Nat Commun 12, 3519, doi:10.1038/s41467-021-23683-y (2021).

48 Chiang, C. Y., Veckman, V., Limmer, K. & David, M. Phospholipase Cgamma-2 and intracellular calcium are required for lipopolysaccharide-induced Toll-like receptor 4 (TLR4) endocytosis and interferon regulatory factor 3 (IRF3) activation. J Biol Chem 287, 3704–3709, doi:10.1074/jbc.C111.328559 (2012).

49 Kholmukhamedov, A., Janecke, R., Choo, H. J. & Jobe, S. M. The mitochondrial calcium uniporter regulates procoagulant platelet formation. J Thromb Haemost 16, 2315–2321, doi:10.1111/jth.14284 (2018).

50 Ajanel, A. et al. Mitochondrial Calcium Uniporter Regulates ITAM-Dependent Platelet Activation. Circ Res 137, 474–492, doi:10.1161/CIRCRESAHA.125.326443 (2025).

51 Ghatge, M. et al. The mitochondrial calcium uniporter regulates calcium dynamics to drive platelet function, bioenergetics, and thrombosis. J Thromb Haemost 24, 716–731, doi:10.1016/j.jtha.2025.10.019 (2026).

52 Cheng, X. et al. Mitochondrial Calcium Overload Drives mtDNA-cGAS-STING Activation via VDAC1 and MCU Upregulation in Periodontitis. Int J Mol Sci 27, doi:10.3390/ijms27104317 (2026).

53 Martyanov, A. A. et al. Effects of bacterial lipopolysaccharides on platelet function: inhibition of weak platelet activation. Sci Rep 10, 12296, doi:10.1038/s41598-020-69173-x (2020).

54 Lopes Pires, M. E., Clarke, S. R., Marcondes, S. & Gibbins, J. M. Lipopolysaccharide potentiates platelet responses via toll-like receptor 4-stimulated Akt-Erk-PLA2 signalling. PLoS One 12, e0186981, doi:10.1371/journal.pone.0186981 (2017).

55 Koessler, J. et al. The Role of Human Platelet Preparation for Toll-Like Receptors 2 and 4 Related Platelet Responsiveness. TH Open 3, e94–e102, doi:10.1055/s-0039-1685495 (2019).

56 Damien, P. et al. NF-kappaB Links TLR1/2 and PAR1 to Soluble Immunomodulator Factor Secretion in Human Platelets. Front Immunol 8, 85, doi:10.3389/fimmu.2017.00085 (2017).

57 Niklaus, M. et al. The involvement of toll-like receptors 2 and 4 in human platelet signalling pathways. Cell Signal 76, 109817, doi:10.1016/j.cellsig.2020.109817 (2020).

58 Suksamai, C. et al. Cyclic GMP-AMP Synthase (cGAS) Deletion Promotes Less Prominent Inflammatory Macrophages and Sepsis Severity in Catheter-Induced Infection and LPS Injection Models. Int J Mol Sci 26, doi:10.3390/ijms26115069 (2025).

59 Chen, Y., Wu, X., Jiang, Z. & Li, X. KAE ameliorates LPS-mediated acute lung injury by inhibiting PANoptosis through the intracellular DNA-cGAS-STING axis. Front Pharmacol 15, 1461931, doi:10.3389/fphar.2024.1461931 (2024).

60 Huang, L. S. et al. mtDNA Activates cGAS Signaling and Suppresses the YAP-Mediated Endothelial Cell Proliferation Program to Promote Inflammatory Injury. Immunity 52, 475–486 e475, doi:10.1016/j.immuni.2020.02.002 (2020).

61 Shimada, K. et al. Oxidized mitochondrial DNA activates the NLRP3 inflammasome during apoptosis. Immunity 36, 401–414, doi:10.1016/j.immuni.2012.01.009 (2012).

62 Kim, J. et al. VDAC oligomers form mitochondrial pores to release mtDNA fragments and promote lupus-like disease. Science 366, 1531–1536, doi:10.1126/science.aav4011 (2019).

63 Yu, C. H. et al. TDP-43 Triggers Mitochondrial DNA Release via mPTP to Activate cGAS/STING in ALS. Cell 183, 636–649 e618, doi:10.1016/j.cell.2020.09.020 (2020).

64 Koupenova, M. et al. Sex differences in platelet toll-like receptors and their association with cardiovascular risk factors. Arterioscler Thromb Vasc Biol 35, 1030–1037, doi:10.1161/ATVBAHA.114.304954 (2015).

65 Koupenova, M. et al. Platelet-TLR7 mediates host survival and platelet count during viral infection in the absence of platelet-dependent thrombosis. Blood 124, 791–802, doi:10.1182/blood-2013-11-536003 (2014).

66 Ahmed, A. et al. Mitochondrial oxidants promote platelet activation and thrombotic susceptibility in prediabetes. J Clin Invest 136, doi:10.1172/JCI195662 (2026).

67 McArthur, K. et al. BAK/BAX macropores facilitate mitochondrial herniation and mtDNA efflux during apoptosis. Science 359, doi:10.1126/science.aao6047 (2018).

68 Howell, N., Nalty, M. S. & Appel, J. A digitonin-based procedure for the isolation of mitochondrial DNA from mammalian cells. Plasmid 16, 77–80, doi:10.1016/0147-619x(86)90083-1 (1986).

69 Dixit, B., Vanhoozer, S., Anti, N. A., O’Connor, M. S. & Boominathan, A. Rapid enrichment of mitochondria from mammalian cell cultures using digitonin. MethodsX 8, 101197, doi:10.1016/j.mex.2020.101197 (2021).

70 Livak, K. J. & Schmittgen, T. D. Analysis of relative gene expression data using real-time quantitative PCR and the 2(-Delta Delta C(T)) Method. Methods 25, 402–408, doi:10.1006/meth.2001.1262 (2001).

